# Accumulation of Dietary and Non-Enzymatic Oxysterols in Preeclamptic Placentas: Insights into Cholesterol Dynamics in Pregnancy Complications

**DOI:** 10.1101/2024.11.13.623499

**Authors:** Lisa Zou, Lisaura Maldonado-Pereira, Elvis Ticiani, Visalakshi Sethuraman, Isoken N. Olomu, Robert Long, Almudena Veiga-Lopez, Ilce G. Medina-Meza

**Affiliations:** Department of Biosystems and Agricultural Engineering, Michigan State University, East Lansing, MI, USA; Department of Pathology, University of Illinois at Chicago; Division of Neonatology, Department of Pediatrics and Human Development, College of Human Medicine, Michigan State University, East Lansing, Michigan, USA; Department of Obstetrics and Gynecology, Sparrow Health System, East Lansing, Michigan, USA

**Keywords:** **dietary** oxysterols, fatty acids, placenta disorders, cholesterol, lipid oxidation

## Abstract

Oxysterols—oxidized derivatives of cholesterol—play key roles in regulating lipid metabolism, oxidative stress, and immune signaling. Despite their biological importance, their presence and function in human placental tissue remain poorly understood, particularly in pregnancy complications such as preeclampsia. Preeclampsia is a hypertensive disorder of pregnancy associated with oxidative stress and altered lipid homeostasis. This study profiled both dietary and non-enzymatic oxysterols in third-trimester human placentas from healthy term pregnancies and those complicated with preeclampsia, delivered at term or preterm and term. Findings indicate significantly higher oxysterol levels in preeclamptic placentas, particularly for 7-ketcholesterol (7-Keto) and 5α,6β-cholestanetriol, (Triol) regardless of gestational age. 5β,6β-epoxycholesterol (O’Callaghan, Woods, & O’Brien, 2001) was higher in term preeclampsia, while total cholesterol levels were significantly lower in preeclamptic placentas. Correlation analyses revealed distinct associations between oxysterols, cholesterol, and total fat content, indicating unique oxidative signatures in preeclamptic pregnancies. These findings contribute to a growing body of evidence suggesting that altered oxysterol profiles may reflect oxidative stress and placental dysfunction. While causality cannot be inferred, oxysterol profiling may hold potential as a complementary biomarker strategy for identifying pregnancies at risk of adverse outcomes.

## Introduction

Preeclampsia is a pregnancy complication defined by the new-onset hypertension and proteinuria (≥ 300 mg/24 h) after 20 weeks of gestation. It is also associated with endothelial dysfunction, hypertension, and end-organ damage. While the etiology of preeclampsia is not clear, evidence suggests that oxidative stress during pregnancy plays a role in both, normal development of the placenta and the pathophysiology of complications such as miscarriage, premature rupture of the membranes (Jauniaux, Poston, & Burton, 2006) and pre-eclampsia (Burton & Yung, 2011).

Physiologically, lipid parameters, including cholesterol, rise during pregnancy (Bartels & O’Donoghue, 2011). However, excessive production of reactive oxygen and nitrogen species (ROS, RNS) can disrupt cellular homeostasis by oxidizing cell membrane lipids, including fatty acids, phospholipids, and cholesterol (Chiarello et al., 2020). Fatty acids play a critical role in supporting cellular structural and metabolic functions, particularly polyunsaturated fatty acids (PUFAs) of the omega-3 or omega-6 series such as linoleic acid (Doskey et al., 2020) and α-linolenic (ALA, omega 3) considered essential fatty acids. Long-chain PUFAs (LCPUFAs) and their metabolites are required at various stages in placental development (Rani, Wadhwani, Chavan-Gautam, & Joshi, 2016). However, an impaired materno-fetal LCPUFA transport is known to occur in pregnancies that are complicated and characterized by an abnormal placental function such as preeclampsia (Hanebutt, Demmelmair, Schiessl, Larqué, & Koletzko, 2008). Cholesterol the major component of mammalian cell membranes, also serves as the precursor of vitamins, bile acids, and steroid hormones - some of which are synthetized by the human placenta. Its abundance in tissue is determined by endogenous synthesis and dietary intake. The placenta plays an active role in regulating cholesterol biosynthesis, metabolism, and transport and exchange between the maternal and fetal compartments (Palinski, 2009).

Oxysterols are cholesterol oxidation products formed due to cholesterol’s susceptibility to oxidation, and differ from the parent cholesterol by the presence of an additional ketone, epoxy, or hydroxyl group in the A or B rings or a hydroxyl group in the side chain (Medina-Meza, Rodriguez-Estrada, Lercker, Soto-Rodriguez, & Garcia, 2011). Oxysterols are biologically active molecules that regulate cholesterol biosynthesis (Bjorkhem, 2002) and act as ligands to G protein-coupled receptors (Vurusaner, Leonarduzzi, Gamba, Poli, & Basaga, 2016) and nuclear receptors (Aye, Waddell, Mark, & Keelan, 2011). They can also affect the immune system by suppressing the production of IgA by B cells (Spann & Glass, 2013).

Oxysterols can be generated either enzymatically or non-enzymatically. Enzymatic pathways are physiologically regulated and primarily mediated by the cytochrome P450 family (Iuliano, 2011; Zmyslowski & Szterk, 2019). In contrast, non-enzymatic oxysterols are formed through cholesterol autooxidation in foods due to storage, heat, and processing. Among the most abundant oxysterols present in foodstuffs are 7-ketocholesterol (7-Keto), 7β-hydroxycholesterol (7β-OH), 5α,6α-epoxide (5,6α-e poxy), 5β,6β-epoxide (L. Maldonado-Pereira, 2021), cholestan-3β,5α,6β-triol (triol), 7α-hydroxycholesterol (7α-OH), and 25-hydroxycholesterol (25-OH-C) (Lisaura Maldonado-Pereira, Carlo Barnaba, & Ilce Gabriela Medina-Meza, 2023). Both altered cholesterol levels and the accumulation of their oxidized metabolites have been confirmed in several pathologies, including atherosclerosis and neurodegenerative diseases (Sottero et al., 2019; Zmyslowski & Szterk, 2019). While oxysterols have been reported in amniotic fluid, maternal blood, and placental tissue with mixed results (Bodzek, Janoszka, Wielkoszynski, Bodzek, & Sieron, 2002), a single study has investigated the presence of the oxysterol 27-hydroxycholesterol in the human placenta (Winkler et al., 2017). However, a comprehensive understanding of the diversity and distribution of oxysterols in placental tissue remains lacking.

An increase in lipid peroxidation products has been reported in maternal serum of healthy pregnancies (Maseki, Nishigaki, Hagihara, Tomoda, & Yagi, 1981). This rise appears to be counterbalanced by enhanced antioxidant activity during pregnancy (Y. P. Wang, Walsh, Guo, & Zhang, 1991). However, in women with preeclampsia, this antioxidant capacity has been reported to be reduced (Basu et al., 2002) potentially leading to an imbalance between oxidation and antioxidation reactions towards oxidative stress, with the placenta as a primary target. Although oxysterols have been implicated in the pathogenesis of several disease states, including atherosclerosis (Zmyslowski & Szterk, 2017) and dementia (Testa et al., 2016), few studies have measured oxysterol concentrations during pregnancy or investigated their relationship with pregnancy outcomes (Rideout et al., 2023). To address this gap, cholesterol, fatty acids, and both dietary oxysterols (Medina-Meza, Vaidya, & Barnaba) and non-enzymatic oxysterols were quantified in human placental tissues collected from healthy pregnancies, as well as from term and preterm preeclampsia cases, and their potential association with preeclampsia was examined.

## Materials and Methods

### Ethics, exclusion criteria, and placenta collection

The institutional review boards at Michigan State University and Sparrow Hospital approved the project. Women admitted for C-section delivery at Sparrow Hospital (Labor & Delivery Service) were asked to participate in the study and provided written informed consent (IRB IRB#17-1359M). Human placentas (Martysiak-Zurowska, Szlagatys-Sidorkiewicz, & Zagierski) were obtained from three groups of patients, as previously described (Sethuraman et al., 2022), 1) healthy term pregnancies (Term; n = 12), 2) term preeclamptic pregnancies (Term PE; n = 8) and 3) pre-term preeclamptic pregnancies (Pre-Term PE; n = 8). Maternal and neonatal anthropometric and clinical data were obtained from the mothers’ and infants’ medical records. Healthy pregnant mothers at term (at ≥ 37 weeks of pregnancy) with no complications and with none of the exclusion criteria (see below) were enrolled as healthy term controls. Mothers with a diagnosis of preeclampsia (at ≥ 36 weeks of pregnancy) were enrolled in the term preeclampsia group, while mothers with a diagnosis of preeclampsia (between 28 to 35 weeks of pregnancy) were enrolled in the preterm preeclampsia group.

Preeclampsia was defined using the American Congress of Obstetricians and Gynecologists criteria (ACOG, 2019): 1) systolic blood pressure of ≥ 140 mmHg or diastolic blood pressure of ≥ 90 mmHg on two times at least 4 h apart after 20 weeks of gestation in a woman with a previously normal blood pressure and 2) proteinuria. Proteinuria was defined as either a 24-h urine protein of >300 mg or protein to creatinine ratio of > 0.3. Subjects with the following conditions were excluded from the study: multiple gestation, clinical chorioamnionitis, chromosomal anomalies, diabetes (gestational, type 1, and type 2), chronic hypertension (pre-existing hypertension), gestational hypertension without a diagnosis of preeclampsia, major cardiovascular diseases, such as myocardial infarction, cholestasis, drug abuse, and smoking. All placentas were processed within 1 h of delivery. A core biopsy was collected ∼2 cm away from the periphery and the umbilical cord attachment and kept frozen until further processing for mRNA and protein expression analyses. Placental tissues were stored at -80 °C for further analysis.

### Materials, Chemicals and Reagents

Methanol was purchased from Sigma-Aldrich (Huang, Elvington, & Randolph, 2015), chloroform from Omni Solv (Burlington, MA), hexane from VWR BDH Chemicals (Batavia, IL), 1-butanol and potassium chloride (KCl) from J. T. Baker (Allentown, PA), diethyl ether from Fisher Chemical (Pittsburgh, PA), and sodium sulfate anhydrous (Na_2_SO_4_) and sodium chloride (Overholt et al.) from VWR. Labeled standards of cholesterol *d-7*, 7α-hydroxycholesterol-*d-7* (7α-OH), 7β-hydroxycholesterol-*d7* (7β-OH), 5,6α-epoxycholesterol-*d7* (5,6α-Epoxy), 5,6β-epoxycholesterol-*d7* (5,6β-Epoxy), 25-hydroxycholestero-*d6* (25-OH) cholestane-3b,5a,6b-triol-*d7* (triol), 7-ketocholesterol-*d7* (7-Keto), 24(S)*-*hydroxycholesterol-d7 *(24-OH* were purchased from Avanti Polar® (Birmingham, AL), 4β-hydroxycholesterol-*d7),* 25(S)-27-hydroxycholesterol (27-OH) from Cayman chemicals® (Ann Arbor MI) and 6 ketocholestanol (6-Keto), 3,6 Dione, 20α-hydroxycholesterol (20-OH), from Steraloids (Newport, R.I.) and purified by using aminopropyl (Liao, Cheng, Li, Lo, & Kang) cartridges (500 mg/3 mL) from Phenomenex® (Torrance, CA).

### Lipid extraction

Before lipid extraction, placenta tissues were thawed, within 4 h and washed three times with phosphate-buffered saline (PBS) to remove any contaminating blood. Then, approximately 200 mg of placenta were placed in a mortar, liquid nitrogen was added, and the sample was gently minced until a fine powder, then transferred to a glass tube with a screwcap. After, 50 μg of 19-hydroxycholesterol and 140 μg of 5α-cholestane plus a mixture of the oxysterols deuterated standards (1 ng/μL each) in ethanol with 0.1% of BHT were added as internal standards for the determination of total cholesterol and oxysterols, respectively. Then, lipid fraction was extracted with 3 mL of hexane/isopropanol (4/1, v/v) with 0.1% of BHT for 3 min, following a sonication for 10 min in cold water. The homogenate was then centrifuged at 2,800 rpm for 10 min at 4 °C. The supernatant was transferred to a new glass tube and the extraction was repeated with the remaining pellet. The supernatants from the two extractions were pooled.

### Total Cholesterol

A quantification method for cholesterol and oxysterols previously developed in our laboratory (Kilvington, Barnaba, Rajasekaran, Laurens, & Gabriela Medina-Meza, 2021; L. Maldonado-Pereira, C. Barnaba, & I. G. Medina-Meza, 2023) was used in this study. Subsequently, cold saponification was done with 1 N KOH solution in methanol left under gentle agitation at a cool temperature and covered from the light overnight. One-tenth of the unsaponifiable matter was subjected to silylation. The sample was mixed with 100 μL of pyridine and 100 μL of Sweeley’s reactive mixture (pyridine/hexamethyldisilazane/trimethylchlorosilane, 10:2:1, v/v/v) at 75 °C for 45 min, dried under nitrogen stream, and dissolved in 1 mL of n-hexane. One microliter of the silylated solution was injected into a gas chromatograph using a fitted with a Zebron™ZB-5HT (Phenomenex, Torrance, CA) capillary column (30 m × 0.25 mm × 0.25 μm). Oven temperature conditions were set up as follows: 260 to 300 °C at 2.5 °C/min, then from 300 to 320 °C at 8 °C/min, and finally hold at 320 °C for 1 min. The injector and detector were both set at 320 °C. Helium was used as the carrier gas at a flow rate of 54.0 mL/min, a split ratio at 1:50, and a pressure constant of 134 kPa.

### Oxysterols Quantification

The other nine-tenths of the sample was used for the quantification of oxysterols. Purification by NH_2_-SPE was performed as previously reported (Lisaura Maldonado-Pereira et al., 2023). The enriched fraction was recovered and evaporated to dryness under a nitrogen stream. Subsequently, the purified fraction was silylated with Sweeley’s reactive mixture (at 75 °C for 45 min), dried under a nitrogen stream, and dissolved in 200 μL of n-hexane. Oxysterols were isolated and identified using the selected ion molecule (SIM) method with a GCMS-QP2010 SE single quadrupole Shimadzu Corporation (Kyoto, Japan). One μL of the silylated sample was injected into a single quad GCMS-QP2010 SE with the following conditions: from 250 to 280 °C at 2 °C/min, hold at 280 °C for 7 min, and from 280 to 315 °C at 1.5 °C/min. Helium was used as a carrier gas (flow rate of 0.37 mL/min); the split ratio was 1:15, and the pressure was 49.2 KPa. The interface temperature for the GC-MS was 320 °C, with the electron multiplier voltage set at 70 kV and the injector at 320 °C. A fused silica column (30 m x 0.25 mm i.d. x 0.25 um thickness) coated with 5% phenyl polysiloxane (Phenomenex Zebron ZB-5) was used.

### Fatty acids methyl esters (Awogbemi, Onuh, & Inambao) profiling

Fatty acid methyl esters (Awogbemi et al.) were prepared according to the transesterification described somewhere else (Maldonado-Pereira, Barnaba, de Los Campos, & Medina-Meza, 2022). One μL of the methylated sample was injected into a gas chromatograph (GC 2010 Shimadzu^®^, Kyoto Japan) equipped with an autosampler (AOC-20s Shimadzu^®^) and a flame ionization detector (Lin et al.). A fused silica DB-WAX (Agilent J&W, Santa Clara, CA) capillary column (30 m × 0.32 mm i.d. × 0.25 μm film thickness) was used to separate the fatty acids. The oven was programmed as follows: 120 °C to 200 °C with a rate of 3 °C/min, then from 200 °C to 240 °C with a rate of 2 °C/min, and held for 2 min. The injector’s pressure was set up at 92.2 kPa. Temperature for both, injector and detector were set at 250 °C. H_2_ was used as carrier gas at 1 mL/min and the split ratio was 5.0. Data acquisition was done on LabSolutions software version 5.106 (Shimadzu^®^, Kyoto, Japan) and peak areas were identified by comparing their retention times to the pure standards (FAME 37 mixture). Fatty acid content were quantified using tridecanoic acid as the internal standard and reported as g of fatty acid per 100 g of sample. Results on fatty acids contents were obtained using the sums: Σ saturated fatty acids (Σ SFA = C8:0 + C10:0 + C11:0 + C12:0 + C13:0 + C14:0 + C15:0 + C16:0 + C17:0 + C18:0 + C20:0 + C21:0 + C22:0 + C23:0 + C24:0); Σ monounsaturated fatty acids (Σ MUFA = C14:1, cis-9 + C15:1, cis-10 + C16:1, cis-9 + C17:1, cis-10 + C18:1, cis-9 + C20:1, cis-11 + C22:1, cis-13 + C24:1, cis-15); Σ polyunsaturated fatty acids (Σ PUFA = C18:2, cis-9,12 + C18:3, cis-6,9,12 + C18:3, cis-9,12,15 + C20:2, cis-11,14 + C20:3, cis-8,11,14 + C20:3, cis-11,14,17 + C20:4, cis-5,8,11,14 + C22:2, cis-13,16 + C20:5, cis-5,8,11,14,17 + C22:6, cis-4,7,10,13,16,19). and Σ trans fatty acids (Σ TFA= C18:1, trans-9 + C18:2, trans-9,12).

### Statistical Analysis

Descriptive statistics were performed, and both mean and confidence intervals (95%) were computed. Since data was not normally distributed, the variable was log-transformed for all analyses. The relationship of individuals’ oxysterols with serum cholesterol and maternal BMI was assessed with Pearson correlation coefficients. Associations between maternal oxysterols and fatty acids and terms were assessed using multivariable linear regression. We considered nulliparity (yes/no), birth weight, gestation age, maternal age, and pre-pregnancy BMI (continuous variables) as potential confounding factors. Confounders with a (*p*-value < 0.05) were considered significant and were further adjusted in the models. Statistical significance was set at a two-sided alpha level of 0.05. When comparing data that did not follow a normal distribution after a density plot analysis and was not homogeneous after a residual plot analysis. Statistical differences between food categories were evaluated by using the non-parametric Kruskal-Wallis ANOVA by Ranks test at *p* < 0.05 significance level. To model the association between metabolite and demographic predictors with pregnancy outcomes (Healthy Term, Term Preeclampsia, and Preterm Preeclampsia), we employed multinomial logistic regression. The analysis utilized a stepwise selection approach for main effects, with BMI and Age included as forced covariates and metabolites entered as stepwise terms using the Forward Stepwise (Likelihood Ratio) method. Covariates were treated as factors to establish hierarchical relationships among predictors. Entry and removal probabilities were set at *p* = 0.05 and *p* = 0.1, respectively, with the likelihood ratio test applied to evaluate variable inclusion and exclusion at each step of the model-building process. All analyses were conducted using SPSS version 29 (IBM®).

## Results and discussion

### Clinical Characteristics

Maternal and neonatal clinical data are presented in **Table 1**. No significant differences were observed among groups in maternal age, height, weight, SROM score, or neonatal birth weight percentiles. Maternal BMI was lower in the Term PE group compared with the Pre-Term PE group. Neonatal birth weight was significantly lower in the Pre-Term PE group compared to the healthy term and Term PE groups. No significant differences were observed in newborn sex distribution across groups.

**Table 1.** Clinical characteristics.

|  | Term (n = 12)<br>≥ 37 weeks |  | Term PE (n = 7)<br>≥ 36 6/7 weeks |  | Pre-Term PE (Laurens)<br>28-35 weeks |  |
| --- | --- | --- | --- | --- | --- | --- |
|  | Mean ± SEM, percent, or N | Range | Mean ± SEM, percent or N | Range | Mean ± SEM, percent, or N | Range |
| Age (years) | 31.8 ± 1.0<sup>a</sup> | 27 - 39 | 25.6 ± 3.0<sup>b</sup> | 19 - 41 | 27.9 ± 1.1<sup>a,b</sup> | 24 - 34 |
| Height (m) | 1.7 ± 0.02 | 1.6 - 1.8 | 1.6 ± 0.03 | 1.5 - 1.7 | 1.6 ± 0.02 | 1.5 - 1.7 |
| Maternal weight (kg) | 87.8 ± 5.6 | 68.5 - 120.2 | 76.4 ± 11.9 | 47.6 - 137.0 | 95.5 ± 8.7 | 68.0 148.8 |
| BMI (kg/m<sup>2</sup>) | 29.6 ± 2.1 | 16.6 - 46.8 | 29.4 ± 4.0 | 20.5 - 50.2 | 36.9 ± 3.2 | 28.3 - 56.3 |
| Birth weight (g) | 3,648 ± 105<sup>a</sup> | 3,090 - 4,240 | 3,081 ± 220<sup>a</sup> | 1,980 - 3,540 | 2,301 ± 228<sup>b</sup> | 1,460 - 3,460 |
| Birth weight percentile | 63.5 ± 6.1 | 30.8 - 99.6 | 43.7 ± 14.0 | 1.8 - 89.7 | 60.2 ± 8.4 | 20.3 - 95.2 |
| Newborn sex (Female/Male) | (4/8) |  | (3/4) |  | (3/6) |  |
| SROM (%) | 0 % |  | 42 % |  | 22 % |  |

Data expressed in mean ± SEM or percent unless otherwise specified. IQR: interquartile range, PE: preeclampsia, Pre-Term PE: preterm preeclampsia, Term PE: term preeclampsia, PTL: preterm labor, SROM: spontaneous rupture of membranes, and T: term control. a ≠ b denotes significant differences between groups.

### Placental fat and fatty acids profile

The fatty acid profiles of human placentas for all three groups are reported in **Table S1.** A total of 31 fatty acids were detected across all placenta samples, whereas heneicosylic acid (C21:0) was only detected in healthy term placentas. No significant differences in the overall fatty acid profile were noted among the three groups.

Surprisingly, elaidic acid (C18:1 trans-9) levels were lower in preeclamptic placentas compared to controls, while the ratio of oleic acid (Lopes et al.) to elaidic acid was significantly higher in preeclamptic placentas. Disturbances in fatty acid metabolism has previously been suggested to contribute to the pathogenesis of preeclampsia (Wadhwani, Patil, & Joshi, 2018). Among these, trans fatty acids, primarily derived from processed foods, have been shown to cross the placenta and accumulate in fetal tissues. Maternal intake of trans fats is reflected not only in fetal circulation but also in breastfed infants (Basak, Vilasagaram, & Duttaroy, 2020; Cohen, Rifas-Shiman, Rimm, Oken, & Gillman, 2011), suggesting a sustained impact of maternal diet on early development. The elevated trans/cis monounsaturated fatty acid ratio observed may indicate early lipid perturbations in the preeclamptic placenta, potentially linked to dietary exposures. Surprisingly, heptadecanoic acid (C17:0), an odd-chain saturated fatty acid considered a potential marker of dairy fat intake (Jenkins, West, & Koulman, 2015), was significantly lower (Term: ∼0.49%; Preterm PE: ∼0.35%; Term PE: ∼0.34%; *p < 0.05*) in preeclamptic placentas, regardless of gestational age. C17:0 has also been associated with a reduced risk of developing multiple sclerosis due to its role in increasing membrane fluidity (Holman et al., 1995). While C15:0 levels (∼0.51–0.75%) did not differ significantly among groups, C17:0 is consistently reported to be more abundant than C15:0 in human adipose tissue (C15:0: ∼ 0.32%, C17:0: ∼0.34%) (Brevik, Veierod, Drevon, & Andersen, 2005), erythrocytes (C15:0: ∼0.28%, C17:0: ∼0.45%) and human hindmilk (C15:0: ∼0.46%, C17:0: ∼0.57%) (Valentine, Hurst, & Schanler, 1994).The ratio of C15:0/C17:0 in human plasma is 1.2 in healthy individuals. Control (1.36) and Pre-Term (1.45) ratios are in agreement with reported values, however, Term-PE differs with a 2.20 ratio. Although odd-chain fatty acids are established markers of dairy fat intake, the observed differences may instead reflect underlying lipid dysregulation associated with pregnancy complications. Specifically, the reduced placental levels of C17:0 in preeclampsia could result from variations in maternal dietary intake, altered endogenous synthesis, or impaired fatty acid transport and metabolism. Further studies are needed to clarify these observations.

Total saturated fatty acids (SFA), monounsaturated fatty acids (MUFA), and PUFAs are significantly reduced (p< 0.05) in placentas with preeclampsia independent of gestational age. These findings align with previous reports of lower maternal LCPUFA levels in early pregnancies (26 - 30 weeks) among women with preeclampsia (Wadhwani et al., 2014), suggesting that altered fatty acid availability may be a feature of the disorder. Potential contributions to these reductions include poor dietary sources, impaired synthesis, or decreased mobilization from maternal stores. Maternal LCPUFAs are critical not only for fetal neurodevelopment but also for proper fetoplacental growth and development. The fetal requirement for fatty acids depends on maternal diet as well as placental transport and metabolism, as the human placenta is permeable to fatty acids and facilitates their transfer to the fetus (Haggarty, 2010) (Wadhwani et al., 2018). Because conversion of precursors such as LA or ALA to LCPUFAs is not sufficient, the mother has to obtain these LCPUFAs directly from the diet to meet fetal demands. This is particularly critical in late gestation, when fetal DHA accumulation reaches 67–75 mg/day to support rapid fetal development. Furthermore, the trophoblast cells require LCPUFA from early pregnancy to carry out various processes associated with optimal placental growth and development. At the same time, the placenta itself relies on an adequate supply of LCPUFAs to support key cellular processes such as trophoblast proliferation, differentiation, oxidative stress regulation, and hormone synthesis—all essential for its development and function (Rani, Chavan-Gautam, Mehendale, Wagh, & Joshi, 2015).

### Cholesterol homeostasis dysregulated with placenta disorders

Essential for fetal development, cholesterol serves as a precursor for steroid hormones and a structural component of cell membranes. In the placenta, cholesterol is taken up from maternal lipoproteins via LDL receptors on the syncytiotrophoblast and transported across apical and basolateral membranes into the fetal circulation. Maternal blood levels of cholesterol, LDL, HDL, and triglycerides are known to rise progressively during the second and third trimesters as part of normal metabolic adaptation to pregnancy(C. Wang et al., 2018; Wild & Feingold, 2023)} In the placenta, cholesterol crosses two barriers (apical and basolateral membrane) to reach the maternal to fetal circulation. Cholesterol travels across the syncytiotrophoblast from the maternal lacunae, which expresses LDL receptors. ABCA and ABGA cholesterol transporters control the efflux of cholesterol into the fetal microcirculation as HDL particles (Palinski, 2009; Woollett, 2005). A significant reduction of total cholesterol in Term-PE placenta in comparison with Term (p < 0.05) was observed. This observation is in contrast with the literature that reports increased levels of cholesterol, triacylglycerides, HDL, and LDL in the second and third trimesters of the pregnancy. However, the majority of reports measure these parameters in blood. These differences may be due that, in current obstetrics settings, free cholesterol is not measured in the placenta after delivery, and there is not a reference range for “healthy” lipid parameters during normal pregnancy. However, far less is known about cholesterol levels within the placental tissue itself; to our knowledge, reference ranges for normal placental lipid content have not been established yet (Bartels & O’Donoghue, 2011). In this study, total cholesterol levels in placental tissue were significantly reduced in Term-PE placentas compared to healthy term placentas (85.14 ± 28.88 µg/mg *vs*. 195.81 ± 134.31 µg/mg; *p < 0.05*; **Table S1** & **Figure 1**). A reduction was also observed in Preterm-PE placentas (126.04 ± 44.49 µg/mg). These findings suggest that placental cholesterol content may be disrupted in preeclampsia, potentially reflecting impaired cholesterol uptake, metabolism, or efflux within the placenta despite elevated circulating cholesterol levels commonly reported in the maternal bloodstream during late pregnancy. Further studies are warranted to clarify how placental lipid handling is altered in preeclampsia and how these changes impact fetal development.

**Figure 1.**
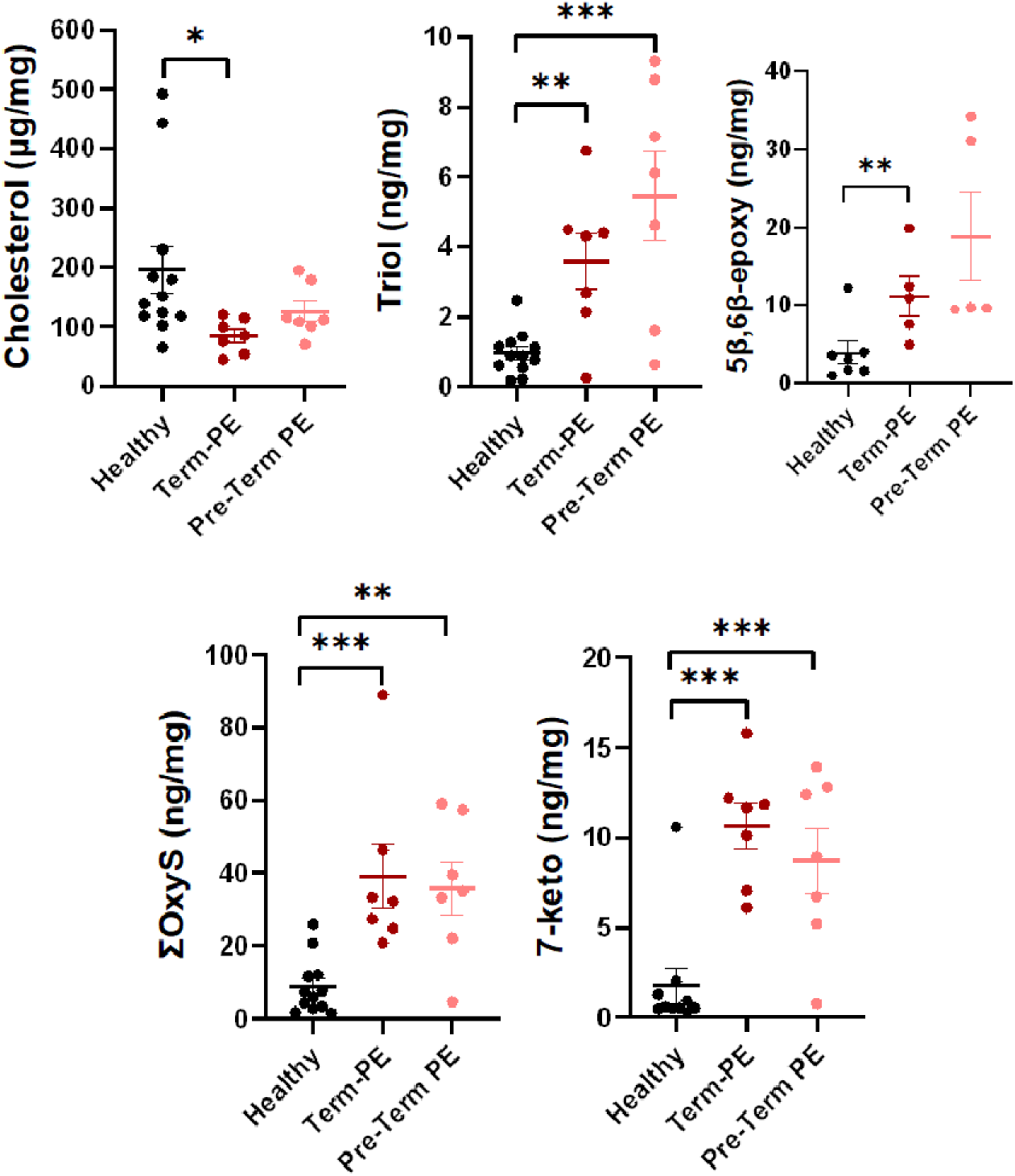
Mean (Abela et al.) of total oxysterols (ΣOxyS), cholesterol, 7-Keto, Triol), 5α,6α-epoxy; 5β,6β-epoxy, 7α-OH and 7β-OH in human placentas derived from age-matched healthy or preeclamptic subjects at term (Pang et al.) or preterm (PE-PTB; <37 weeks GA). *P<0.05, **P<0.005, ***P<0.0005 by ANOVA.

### Presence of oxysterols in the human placenta

In this study, we report for the first time the presence of 11 oxysterols in human placentas with preeclampsia at different terms **(Table 2** & **Figure 1)**. Six oxysterols (7-OH isomers, 5,6-isomers, 7-keto, and Triol) were detected in all human placentas (Pustjens, Boerrigter-Eenling, Koot, Rozijn, & van Ruth), representing up to 70% of the total oxysterol content in human placentas. The ratio between individual oxysterols and the total oxysterol (OxyS) content in placental tissue is shown in **Table 3**. The sum of all oxysterols was two-fold higher in preeclamptic placentas, independent of gestational age (Term: 8.76 ng/mg Term-PE: 39.12 ng/mg; Preterm-PE: 35.87 ng/mg; *p < 0.001*) **(Figure 1)**. To note, when analyses were conducted including maternal body weight as a covariate, all significant differences in oxysterols among groups remained.

**Table 2.** Percent of oxysterol abundance in human placentas.

| Oxysterol | ID | Abundance in human placenta (%) |
| --- | --- | --- |
| 7α-hydroxycholesterol | 7α-OH | 92.9 |
| 7β-hydroxycholesterol | 7β-OH | 92.9 |
| cholesterol 3β,5α,6β-triol | Triol | 92.9 |
| 7-ketocholesterol | 7-Keto | 85.7 |
| cholesterol 5α,6α-epoxide | 5α,6α-epoxy | 64.3 |
| cholesterol 5β,6β-epoxide | 5β,6β-epoxy | 60.7 |
| 25-hydroxycholesterol | 25-OH | 25.0 |
| 22-hydroxycholesterol | 22R-OH | 21.4 |
| 6-ketocholestanol | 6-Keto | 14.3 |
| 20S-hydroxycholesterol | 20-OH | 10.7 |
| 24-hydroxycholesterol | 24-OH | 7.1 |

**Table 3.** The ratio between individual oxysterols and the total oxysterol (OxyS) content in placental tissue.

| Ratio | Control (n=12) |  | Pre-Term PE (n=9) |  | Term- PE (n=7) |  | *p-value* |
| --- | --- | --- | --- | --- | --- | --- | --- |
|  | mean | SD | mean | SD | mean | SD |  |
| Total OxyS (ng/mg) | 8.76 | 7.74 | 35.87 | 19.09 | 39.12 | 23.42 | **0.001** |
| 7α-OH/OxyS | 0.21 | 0.06 | 0.10 | 0.13 | 0.04 | 0.02 | **0.001** |
| 7β-OH/OxyS | 0.26 | 0.10 | 0.09 | 0.10 | 0.04 | 0.02 | **0.001** |
| 5,6β/OxyS | 0.28 | 0.15 | 0.39 | 0.16 | 0.33 | 0.12 | |
| 5,6α/OxyS | 0.03 | 0.01 | 0.07 | 0.04 | 0.05 | 0.03 | |
| Triol /OxyS | 0.19 | 0.14 | 0.16 | 0.08 | 0.13 | 0.10 |  |
| 7-keto/OxyS | 0.15 | 0.13 | 0.27 | 0.17 | 0.33 | 0.16 |  |
| 25-OH/OxyS | 0.04 |  | 0.17 |  | 0.13 | 0.10 |  |
| 7α OH/7β OH | 0.86 | 0.25 | 1.15 | 0.79 | 1.15 | 0.26 | |
| 5,6α Epoxy/5β Epoxy | 9.84 | 6.24 | 5.97 | 2.07 | 7.96 | 6.03 | |

Similarly to reported oxysterol serum levels, 7-keto, 5β,6 β-epoxy, and Triol were increased and positively correlated with preeclampsia independent of gestational age (term or preterm). These oxysterols are formed non-enzymatically, often generated through ROS-induce lipid peroxidation or derived from dietary sources. Notably, 5β,6β-epoxy was significantly higher in Term, but not Pre-Term preeclamptic placentas. This is consistent with previous finding in maternal serum, where women with pregnancy-induced hypertension had significantly higher levels of 5,6-epoxide isomers, alongside an overall increase in total oxysterols (p=0.042). Our findings align with Sevanian et al., (1994) who reported that 5β,6β epoxycholesterol rather than the 5α,6α isomer was the most abundant epoxide in human plasma. In contrast, other studies have identified side chains oxysterols, such as 27-OH (144.5 ng/mL) and (24-OH (84.9 ng/mL) as predominant in blood during mid-gestation, followed by 7α-OH (47.4 ng/mL) and 24-OH (18.4 ng/mL). However, these studies did not report epoxide levels (A. Dickson et al., 2023) More recently Dickson et al. (A. L. Dickson, Yutuc, Thornton, Wang, & Griffiths, 2022) found that C-7 derived oxysterols were the most abundant in plasma in the following order: 7β-OH > 7α-OH > 26-OH > 24-OH. These discrepancies may reflect differences in study design, detection methods, gestational timing, or nutritional status, and underscore highlight the need for harmonized quantification methods and dietary assessment to better distinguish endogenous from exogenous sources of oxysterols.

The presence of 20-OH in placental tissue was confirmed, consistent with previous findings (A. L. Dickson et al., 2022). Although infrequently detected in clinical studies, 20-OH is biologically active and has been shown to inhibit the activation of SREBP-2 (Nohturfft, Yabe, Goldstein, Brown, & Espenshade, 2000), a key transcription factor regulating cholesterol biosynthesis, potentially through binding to INSIG. Along with side-chain oxysterols (22R-OH and 20R, 22R-dihydroxycholesterol), 20-OH also acts as a ligand for liver X receptors (Vieira et al.) In contrast, the most abundant oxysterols associated with dietary intake, such as 5α,6α-epoxy, 7α-OH, and 7β-OH did not significantly differ between healthy term and preeclamptic placentas.

### Associations of oxysterols with placental fat, cholesterol and FFA

Correlation analyses revealed distinct patterns between placental oxysterol concentrations and total cholesterol levels **(Figure 2)**. This included a positive correlation between 7α-OH and 7β-OH (r^2^: 0.741; *p < 0.01* and 0.548; *p < 0.0001*) and between 5α,6α-Epoxy and 5β,6β-Epoxy (r^2^: 0.925;p< 0.01 and 0.855; *p < 0.0001*). Notably, cholesterol concentrations were positively correlated with 7α-OH and 7β-OH and negatively correlated with Triol and 7-Keto (**Figure S1,** Supplemental material). **Figure 3** shows the ratio of total placental fat to cholesterol or oxysterols across groups. Compared to healthy controls, both PE-Term and PE-Preterm placentas exhibited higher fat-to-cholesterol and fat-to-oxysterol ratios, particularly for 7-keto, Triol, and 5β,6β-epoxy. These shifts suggest altered lipid partitioning and disproportionate oxysterol accumulation relative to overall fat content in preeclamptic placentas.

**Figure 2.**
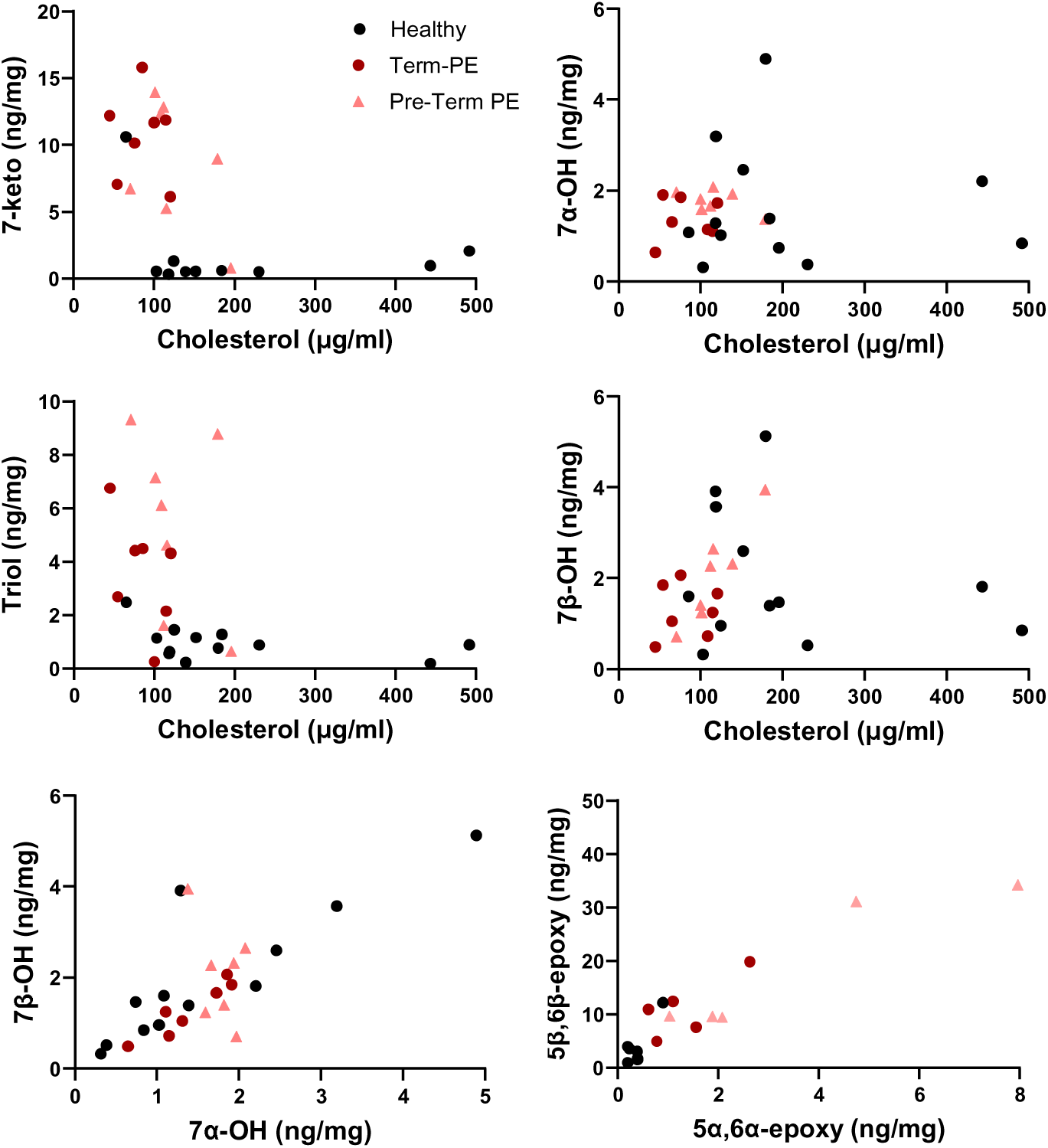
Correlation plots between oxysterols (OxyS) and cholesterol in human placentas derived from age-matched healthy or preeclamptic subjects at term (Pang et al.) or preterm (PE-PTB; <37 weeks GA).7-Keto, Triol, 5β,6β-Epoxy7α-OH and 7β-OH.

**Figure 3.**
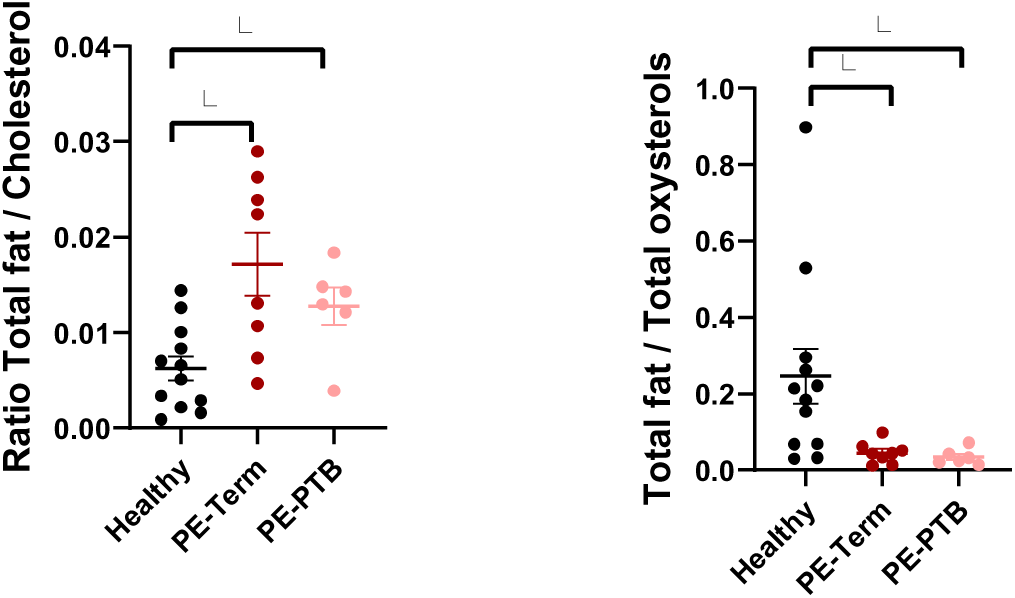
Ratio of total placental fat and cholesterol or oxysterols in human placentas derived from age-matched healthy or preeclamptic subjects at term (Pang et al.) or preterm (PE-PTB; <37 weeks GA). *P<0.05 by ANOVA.

Ratios of SFA, MUFA, and PUFA to both Oxys and cholesterol were significantly altered in preeclamptic placentas compared to healthy controls **(Figure 4)**. SFA/Oxys, MUFA/Oxys, and PUFA/Oxys ratios were significantly reduced in both Term-PE and Pre-Term PE groups relative to Healthy, with the most pronounced decrease observed in the PUFA/Oxys ratio (*p < 0.001*). In contrast, SFA/Cholesterol and PUFA/Cholesterol ratios were significantly elevated in PE groups, particularly in Term-PE placentas (*p < 0.01* to *p < 0.001*). MUFA/cholesterol ratios, however, showed no significant differences among groups. These findings suggest a disproportionate accumulation of cholesterol and oxidized lipid species relative to fatty acids in preeclamptic placentas.

**Figure 4.**
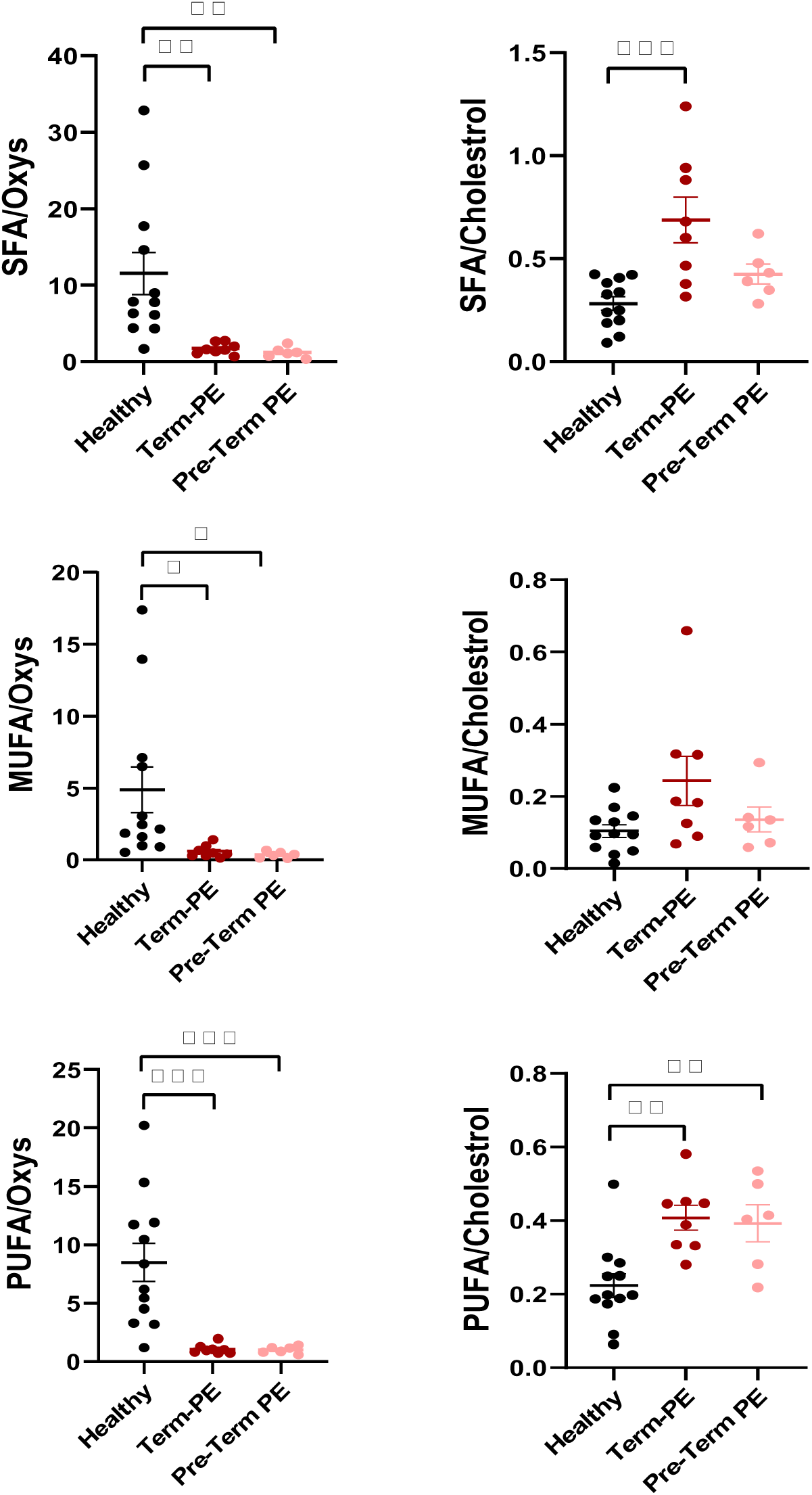
Correlation plots between saturated (SFA), monounsaturated (MUFA), and polyunsaturated fatty acids (PUFAs) with cholesterol and OxyS p<0.05 in human placentas derived from age-matched healthy or preeclamptic subjects at term (Pang et al.) or preterm (PE-PTB; <37 weeks GA). *P<0.05, **P<0.005, ***P<0.0005 by ANOVA.

Multinomial logistic regression (Ezeamama et al.) was performed to model the association between cholesterol metabolites and fatty acids with pregnancy outcomes, using *Term* as the reference category. MLR allows for the analysis of relationships between multiple predictors and a categorical dependent variable with more than two levels (Li, Rysavy, Bobashev, & Das, 2024). An initial model including age and BMI showed no improvement in model performance, so these variables were excluded from the final analyses. Stepwise selection identified 5,6α-epoxycholesterol and its metabolite Triol as significant predictors (*p < 0.001*). However, due to high collinearity between these variables only Triol, that demonstrated a stronger and more reliable association with pregnancy outcomes, was retained to ensure model stability (**Table 4**). The model demonstrated strong predictive power to distinguish Pre-Term PE from healthy term pregnancies (*p = 0.016*), but lacked sufficient predictive power for detecting Term-PE (p = 0.215). The classification accuracy was highest for Pre-Term PE pregnancies (91.7%) followed by healthy term (71.4%) and Term-PE (57.1%), yielding an overall classification accuracy of 76.9%. This reduced performance in the Term-PE category is likely attributable to limited sample size. **Figure 5** presents a scatter plot of the estimated classification probabilities for predicted *vs*. actual categories, illustrating the degree of alignment between predicted probabilities and true outcomes.

**Figure 5.**
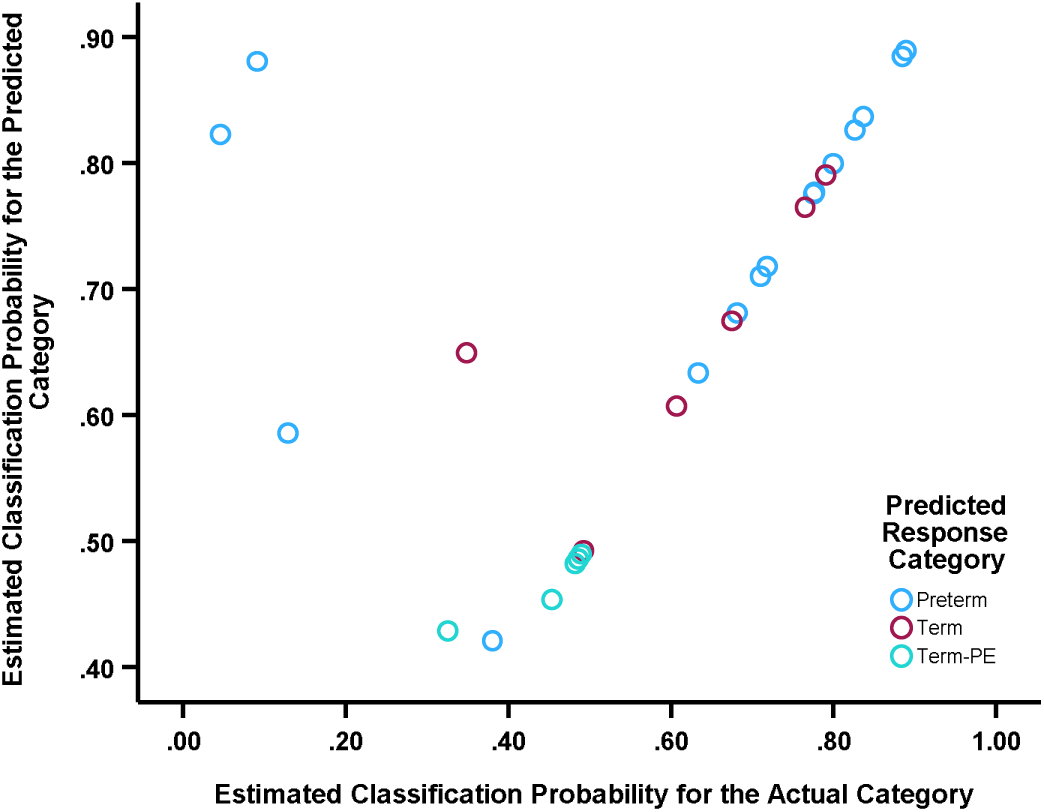
Scatter plot of estimated Classification probabilities for predicted *vs*. actual categories. The plot shows the model’s predicted probabilities against the actual probabilities for each observation, highlighting the alignment between predicted and true category classifications.

**Table 4.** Parameter estimates from multinomial logistic regression model for predicting pregnancy outcomes.

| Terms<sup>a</sup> |  | B | Std. Error | Wald | df | Sig. | Exp(B) | 95% Confidence Interval for Exp(B) |  |
| --- | --- | --- | --- | --- | --- | --- | --- | --- | --- |
|  |  |  |  |  |  |  |  | Lower Bound | Upper Bound |
| **Preterm** | Intercept | 3.814 | 1.307 | 8.512 | 1 | **0.004** |  |  |  |
|  | Triol | -1.425 | 0.590 | 5.830 | 1 | **0.016** | 0.240 | 0.076 | 0.765 |
| **Term-PE** | Intercept | 1.241 | 1.145 | 1.176 | 1 | 0.278 |  |  |  |
|  | Triol | -0.276 | 0.223 | 1.535 | 1 | 0.215 | 0.759 | 0.490 | 1.174 |

<sup>a</sup>The reference category is: Term.

Triol has been studied for its potential involvement in inflammatory and oxidative stress pathways across various health conditions. In human cells, triol is derived from the hydrolyzation of 5,6-epoxycholesterol isomers by cholesterol-5,6-epoxy hydrolase (Poirot & Silvente-Poirot, 2013). It serves as a precursor of two highly bioactive sterols implicated in tumorigenesis, dendrogenin A and oncosterone (Dalenc, Poirot, & Silvente-Poirot, 2015; de Medina et al., 2021), and it has been proposed as a marker for Niemann-Pick Type C disease (Reunert et al., 2015) and subclinical-hypothyroidism (Unluturk et al., 2021). However, further investigations are needed to determine whether elevated Triol levels contribute causally to pregnancy complications. The current study is limited by a small sample size and thus further research will be needed to evaluate the utility of Triol as a biomarker for pregnancy complications such as preeclampsia.

## Conclusion

This study presents novel data on the presence of both dietary and enzymatically generated oxysterols in third-trimester human placental tissue. Notable differences in oxysterol concentrations were observed across pregnancy outcomes, particularly in relation to preeclampsia and preterm birth. However, interpretation of these findings is limited by the relatively small sample size, and further studies are needed to confirm the utility of oxysterols as biomarkers of pregnancy complications.

Unlike cholesterol, which is commonly monitored in non-pregnant adults, cholesterol and oxysterols are not routinely assessed during pregnancy, and standardized reference ranges do not currently exist. Emerging evidence suggests that elevated maternal cholesterol and oxysterol levels during pregnancy are associated with an increased risk of preeclampsia (PE) and preterm birth. However, further research is needed to establish physiologically normal ranges, clarify the role of maternal hypercholesterolemia, and determine whether dietary or pharmacological interventions can mitigate these risks. Although this study does not provide causal evidence for a direct role of oxysterols in the pathogenesis of preeclampsia, the findings add to a growing body of literature indicating that placental lipid metabolism and oxysterol accumulation may be disrupted in pregnancy disorders. Ultimately, profiling oxysterols in the placenta could become a valuable tool for identifying pregnancies at risk for complications, but this potential must be validated through larger, prospective studies.

## Acknowledgments

We thank the Pathology Department at Sparrow Hospital for their help in sample collection. This study was funded by the USDA, Hatch project MICL02526 and Michigan State University start-up funding to I.G.M.M. and partially funded by the National Institutes of Environmental Health Sciences 1R01ES035691 to A.V.-L.)

## Author contributions

**L. Zou:** formal analysis and data curation; **L. Maldonado-Pereira:** formal analysis and data curation; **Elvis Ticiani:** data analyses; **Visalakshi Sethuraman:** patient recruitment, sample collection, data abstraction; **Isoken Nicholas Olomu:** patient recruitment, data abstraction; **Robert Long:** patient recruitment; **A. Veiga-Lopez:** Conceptualization, funding acquisition, investigation, data curation, formal analysis, writing – original draft, writing – review and editing; **I. G. Medina-Meza:** Conceptualization, investigation, funding acquisition, writing – original draft, methodology, data curation, supervision and formal analysis, review and editing.

## Ethical approval

Ethics approval was not required for this research.

## Data availability statement

All data are included in the manuscript.

## SUPPLEMENTAL INFORMATION

**Figure S1.**
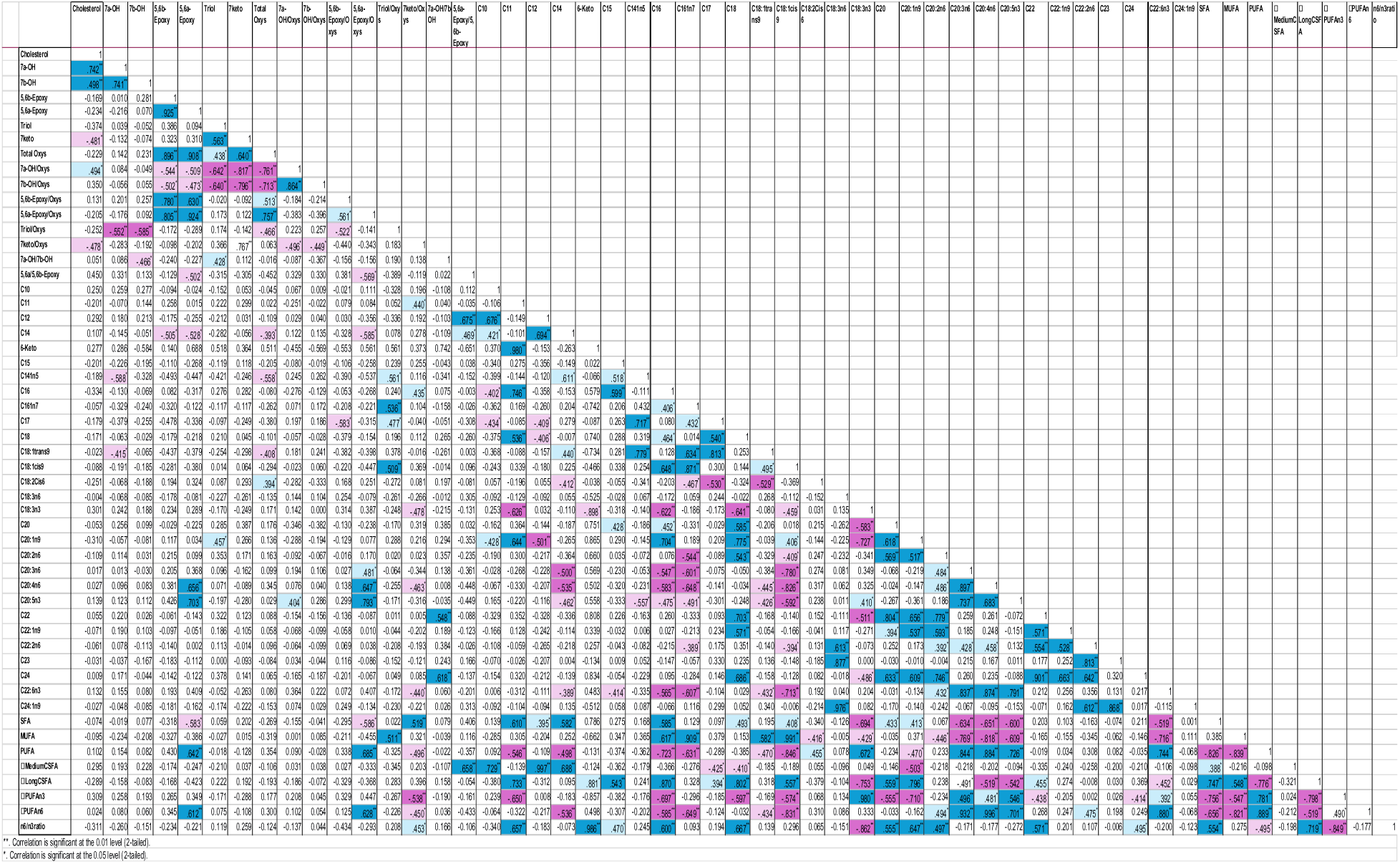
Placental lipid metabolites correlations.

**Table S1.**
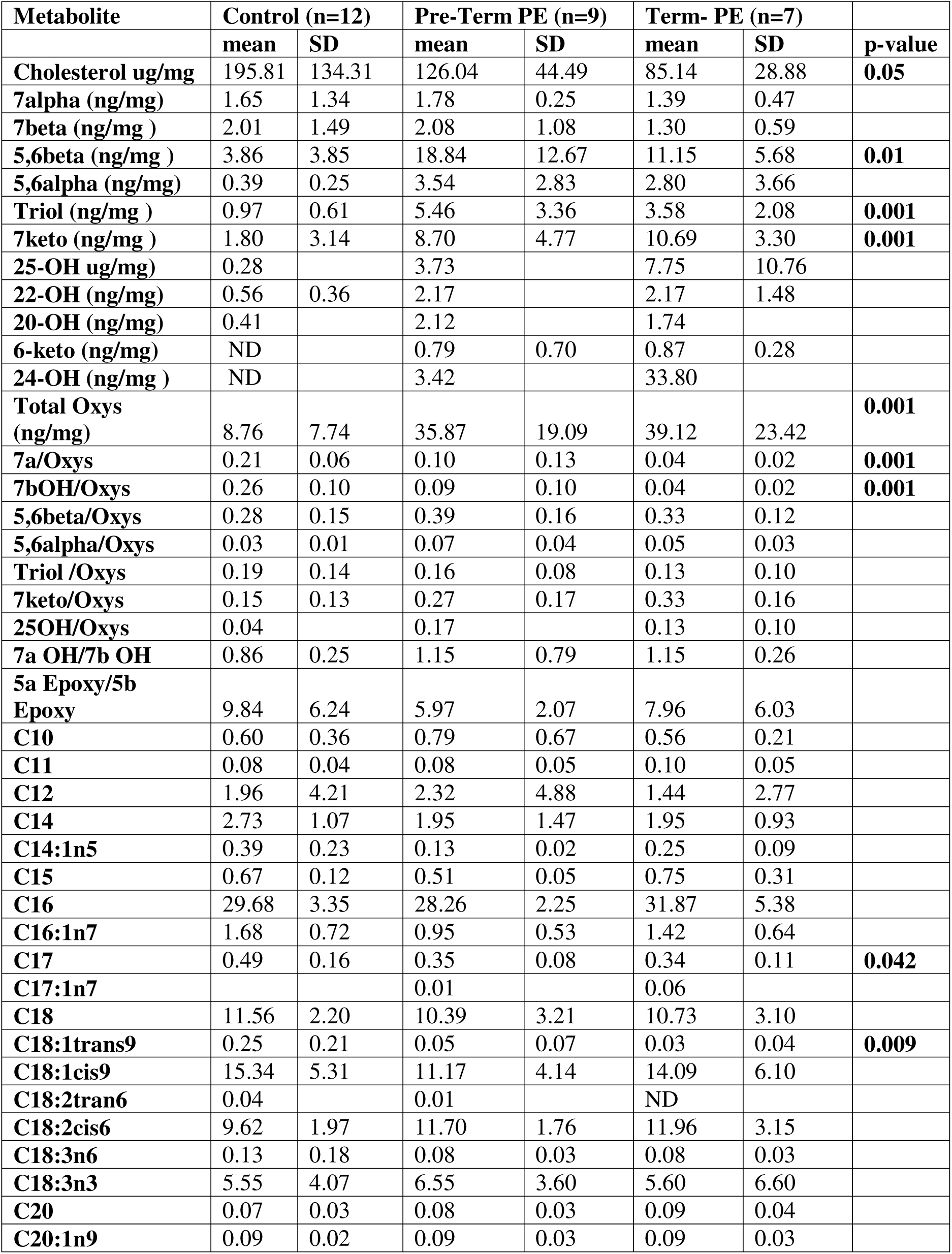
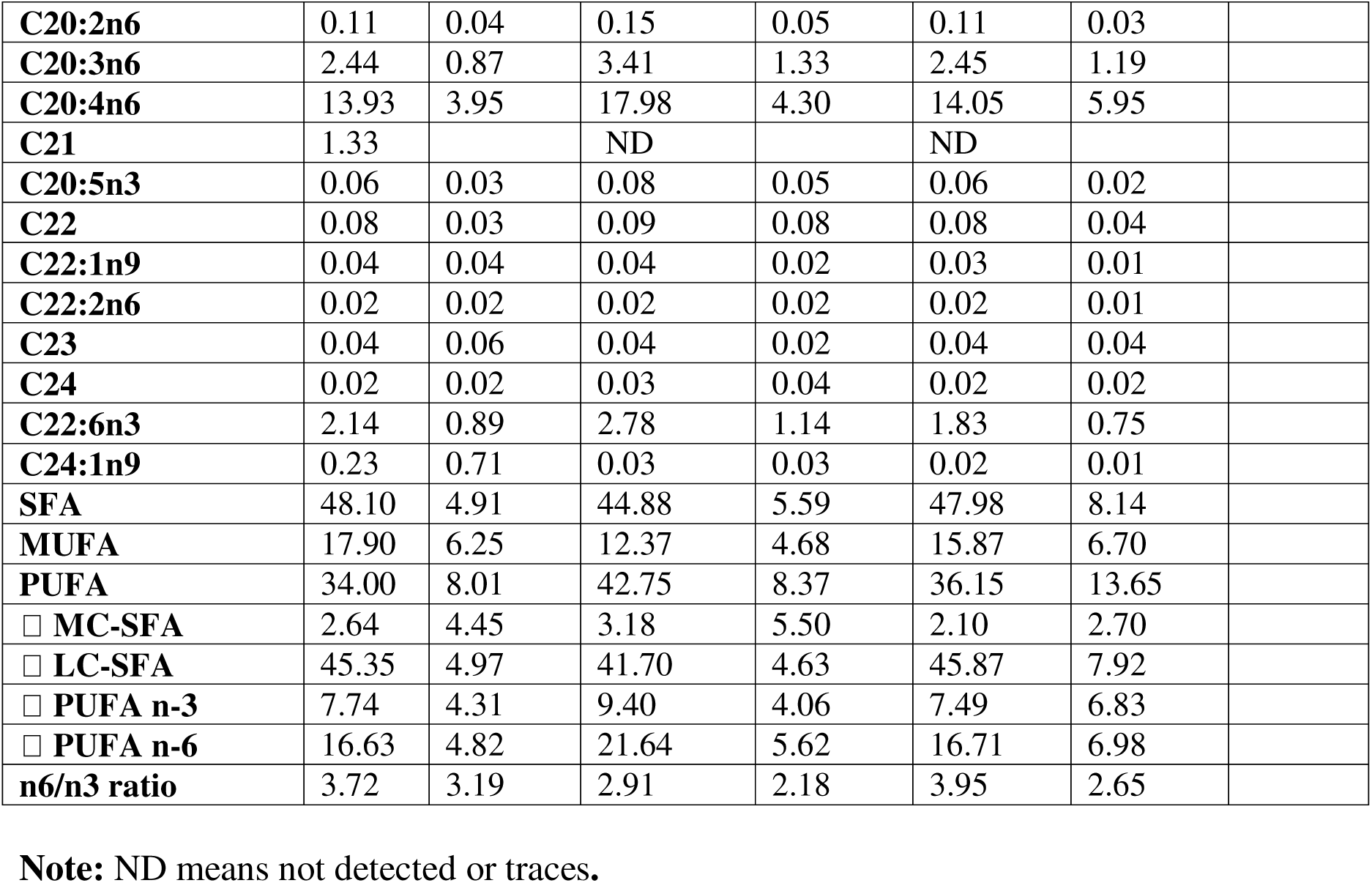
Oxysterols and fatty acids profile in human placentas.

